# Dynamics of Notch-Dependent Transcriptional Bursting in its Native Context

**DOI:** 10.1101/496638

**Authors:** ChangHwan Lee, Heaji Shin, Judith Kimble

**Affiliations:** Department of Biochemistry, University of Wisconsin-Madison, Madison, Wisconsin 53706, USA; Howard Hughes Medical Institute, University of Wisconsin-Madison, Madison, Wisconsin 53706, USA; Current address: The David H. Koch Institute of Integrative Cancer Research, Massachusetts Institute of Technology, Cambridge, Massachusetts 02139, USA

**Keywords:** Notch signaling, MS2 system, Gradient, Transcriptional bursting, Burst duration, Stochasticity, Stem cells, *C. elegans* germline, *sygl-1*, live imaging

## Abstract

Transcription is well known to be inherently stochastic and episodic, but the regulation of transcriptional dynamics is not well understood. Here we analyze how Notch signaling modulates transcriptional bursting during animal development. Our focus is Notch regulation of transcription in germline stem cells of the nematode *C. elegans*. Using the MS2 system to visualize nascent transcripts and live imaging to record dynamics, we analyze bursting as a function of position within the intact animal. We find that Notch-dependent transcriptional activation is indeed “bursty”; that wild-type Notch modulates burst duration (ON-time) rather than duration of pauses between bursts (OFF-time) or mean burst intensity; and that a mutant Notch receptor, which is compromised for assembly into the Notch transcription factor complex, primarily modifies burst size (duration x intensity). To our knowledge, this work is the first to visualize regulation of metazoan transcriptional bursting by a canonical signaling pathway in its native context.

**HIGHLIGHTS:**

- Notch-dependent transcriptional bursts are spatially graded across the stem cell pool
- Burst duration is the key determinant of Notch-dependent transcriptional probability
- Notch NICD strength influences both burst duration and intensity
- Notch dynamics are largely stochastic for consecutive bursts at the same chromosomal locus

## INTRODUCTION

Transcriptional dynamics have entered a new era (Liu and Tjian, 2018; Nicolas et al., 2017). Classical studies discovered dynamic transcriptional responses to metabolites (e.g. Jacob and Monod, 1961) as well as dynamic spatio-temporal transcriptional patterns during development (e.g. De Robertis et al., 2000; McGinnis and Krumlauf, 1992). Yet the past decade of now neo-classical studies revealed that nascent transcripts are generated in dynamic “bursts” in virtually all cells from bacteria to humans (Chubb et al., 2006; Golding et al., 2005; Raj et al., 2006). This universal phenomenon raises new and exciting questions about how bursting is modulated by transcriptional regulators. Although progress has been made on this front (Corrigan and Chubb, 2014; Kafri et al., 2016; Larson et al., 2013; Molina et al., 2013; Senecal et al., 2014), most studies have relied on indirect measures (e.g. luminescence of reporter protein) and have been conducted in non-native systems (e.g. tissue culture cells). Only a few pioneering studies have visualized transcriptional bursting in the native context of *Drosophila* embryos (Bothma et al., 2014; Fukaya et al., 2016; Lucas et al., 2013). A major gap in our understanding is how intercellular signaling and dedicated transcriptional regulators modulate bursting in an intact metazoan as they guide development in its native context.

Here we address this gap by analyzing the dynamics of the transcriptional response to Notch signaling – in an intact animal as Notch maintains stem cells within their niche. Notch signaling is central to many aspects of development across metazoan phylogeny, and when aberrant can cause human disease (Artavanis-Tsakonas et al., 1999; Kopan and Ilagan, 2009). The backbone of the Notch molecular mechanism is conserved across animal phylogeny (Bray, 2016; Kovall et al., 2017). Briefly, the binding of Notch ligands expressed on the surface of the signaling cell to Notch receptors expressed on the surface of the receiving cell triggers receptor cleavage. The liberated Notch intracellular domain (NICD) then enters the nucleus and assembles into a complex to activate transcription of Notch-dependent genes. Many studies have analyzed the Notch transcriptional response *in vivo* (e.g. Hoyle and Ish-Horowicz, 2013; Ilagan et al., 2011; Imayoshi et al., 2013; Jenkins et al., 2015; Kershner et al., 2014; Shimojo et al., 2008), but only one smFISH study had sufficient resolution to reveal its probabilistic nature (Lee et al., 2016). Notch is thus a prime candidate for understanding how a canonical signaling pathway regulates the dynamics of transcriptional bursting. A recent study reported that different Notch ligands elicit responses with distinct dynamics, but this was done in cultured cells and did not directly assess transcriptional bursting (Nandagopal et al., 2018). An approach that directly assesses Notch-dependent transcriptional dynamics in its *in vivo* context is therefore timely.

We focus our study on GLP-1/Notch signaling in the *C. elegans* gonad (Figures 1A and 1B) for several reasons. First, its biological context is both important and conserved. Notch maintains stem and progenitor cells from nematodes to vertebrates (Austin and Kimble, 1987; Duncan et al., 2005; Gaiano and Fishell, 2002; van Es et al., 2005). In the nematode, GLP-1/Notch signaling is the major regulator that maintains germline stem cells (GSCs) (Austin and Kimble, 1987). Second, the tissue architecture is simple, well defined, and accessible to imaging within an intact transparent animal. Notch ligands are expressed in a well-defined mesenchymal cell that provides the niche (Henderson et al., 1994; Nadarajan et al., 2009; Tax et al., 1994), whereas GLP-1/Notch receptors are expressed in GSCs (Crittenden et al., 1994). Third, the key downstream genes are known. GLP-1/Notch activates transcription of *sygl-1* and *lst-1*, which are themselves crucial for stem cell maintenance (Kershner et al., 2014; Shin et al., 2017). Indeed, GLP-1/Notch and its key targets maintain a pool of ~50 germ cells in a naïve stem cell-like state (Cinquin et al., 2010). Fourth, signaling is sustained throughout the life of the animal to continuously maintain stem cells (Austin and Kimble, 1987). This system therefore provides an exceptionally tractable platform to analyze how GLP-1/Notch regulates transcriptional bursting.

**Figure 1.**
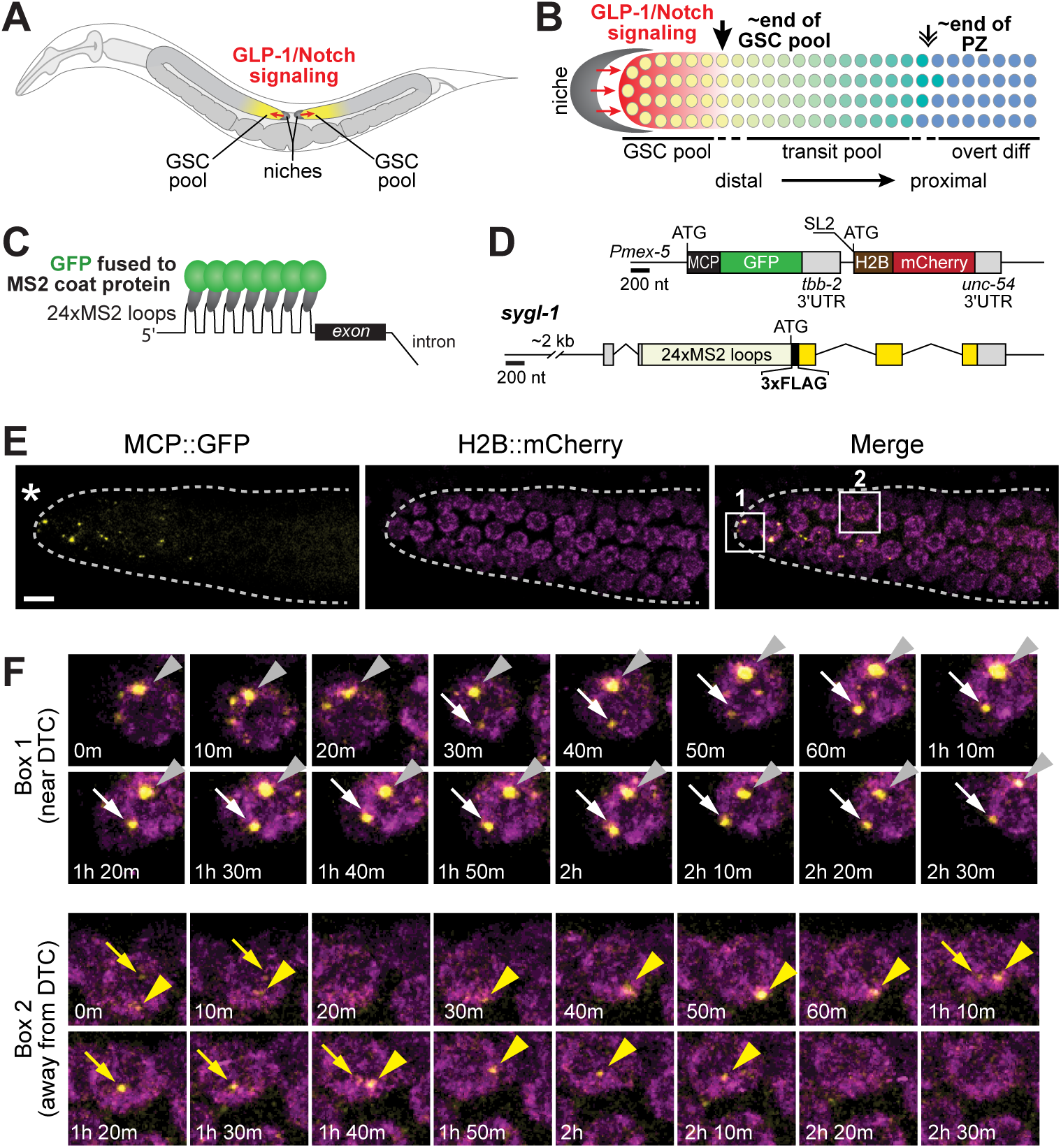
Live imaging of Notch-dependent transcriptional activation. (A) Diagram of *C. elegans* adult. GLP-1/Notch signaling (red) from the niche (dark grey) maintains a pool of germline stem cells (GSCs, yellow) at the distal end of U-shaped gonadal arms (light grey). (B) Diagram of distal gonad with GLP-1/Notch sending and receiving cells. The single-celled niche (dark grey) expresses Notch ligands; the GLP-1/Notch receptor is expressed in naive germ cells in the GSC pool (yellow) as well as in transit cells as they move along the distal to proximal axis towards overt differentiation (green to blue). These naïve and transit pools make up the progenitor zone. The GSC pool ends 6-8 germ cell diameters (gcd) on average from the niche (downward black arrow) while the progenitor zone (PZ) ends 19-22 gcd from the niche (double arrow). smFISH revealed a gradient of GLP-1/Notch-dependent active transcription sites across the GSC pool (graded red) (Lee et al., 2016). (C) MS2 system. MS2 coat protein (MCP) fused to GFP binds to MS2 loops in RNA. Note that MCP binds as a dimer and the number of MS2 loops is actually 24, neither of which are depicted for simplicity. (D) Transgenes for MS2 system used in this work. Exons are boxes, introns are connecting lines. *Above*, operon encoding MCP::superfolder GFP to visualize *sygl-1* transcripts and H2B::mCherry to mark nuclei. The operon is driven by the *mex-5* germline promoter, but trans-spliced to produce two transcripts. SL2, trans-spliced leader. *Below, sygl-1* gene with 24 MS2 loops inserted into the *sygl-1* 5’UTR, just before start codon. Exons include untranslated (grey) and coding regions (blue) (E) Visualization of Notch-dependent *sygl-1* nascent transcripts in the distal gonad of a living adult (24 h post mid-L4 stage), using the MS2 system and confocal imaging. Asterisk marks the distal end of the gonad, where the niche resides; dashed lines mark the gonadal outlines. Scale bar, 5 μm. (F) Montages of boxed regions (1 or 2) in E. Each arrow or arrowhead traces a single MCP::GFP dot over time, as seen in the time-lapse movie (Movie 1). Time points are shown in each image.

The stage was set for this current work by a single-molecule FISH (smFISH) analysis of the Notch transcriptional response at endogenous *sygl-1* and *lst-1* loci in GSCs (Lee et al., 2016). Both nascent transcripts and mature mRNAs were visualized at high resolution and quantitated as a function of cell position within the GSC pool. The generation of active transcription sites (ATS) at both *sygl-1* and *lst-1* was Notch-dependent and stochastic, as predicted; yet the probability of their activation was unexpectedly graded across the pool and that gradation was found to reflect a gradient in Notch signaling strength (Figure 1B). The *sygl-1* and *lst-1* mRNAs and proteins, by contrast, were expressed more uniformly (Lee et al., 2016; Shin et al., 2017), highlighting the need to focus specifically on nascent transcripts to analyze the graded Notch effect on transcriptional bursting.

The *C. elegans* gonad is thus poised to understand how Notch modulates transcriptional bursting in a native context. In this work, we focus on live imaging of *sygl-1* nascent transcripts to confirm the existence of transcriptional bursting, and to quantitate burst features as a function of position within the stem cell pool. Arguably our most important conclusion is that wild-type Notch signaling modulates or “tunes” the duration of active transcriptional bursts, but has little or no effect on duration of the inactive pauses between bursts or burst intensity. This result contrasts with conclusions of other studies, mostly in tissue culture, which highlight burst frequency as the primary target of regulation (see Discussion).

## RESULTS

### Live imaging of Notch-dependent transcriptional activation

To visualize the dynamics of Notch-dependent transcription, we implemented the MS2 system in the *C. elegans* germline. This system relies on a high affinity interaction between MS2 coat protein (MCP) and MS2 RNA loops to bring GFP to transcripts (Figure 1C) (Bertrand et al., 1998; Larson et al., 2009). We used two integrated transgenes to express the system in germ cells (Figure 1D). The first is an operon that employs a strong germline promoter, *mex-5*, to drive expression of two proteins, MCP fused to superfolder GFP (MCP::GFP hereafter) to detect nascent transcripts and histone subunit H2B fused to mCherry to mark nuclei. The second carries a Notch target gene, *sygl-1*, plus 24 MS2 loops inserted into its 5’ UTR. Without MS2 loops, this transgene rescues a *sygl-1* null mutant, but with MS2 loops, it makes no SYGL-1 protein (Figure S1A). Therefore, overall SYGL-1 abundance is likely not affected.

Our MS2 system allows visualization of *sygl-1* nascent transcripts in living animals (Figures 1E and 1F). To image them over time, we immobilized intact animals on a microscope slide, using microbeads and serotonin as previously reported for other *C. elegans* live imaging (Kim et al., 2013; Rog and Dernburg, 2015). This treatment impeded body movement, but not pharyngeal pumping or egg laying (see Methods). Moreover, in the distal third (1-7 gcd) of the progenitor zone (PZ), our region of interest for this study, this treatment did not affect either the rate of germ cell movement along the distal-proximal axis or the frequency of germ cell divisions (see Methods), both consistent with previous studies (Crittenden et al., 2006; Gerhold et al., 2015; Rosu and Cohen-Fix, 2017). Once immobilized, we used a confocal microscope equipped with a temperature-controlled stage, set at 20°C, to image the distal two-thirds of the PZ in live animals at 5-minute intervals for extended periods (three to nine hours). MCP::GFP dots and H2B::mCherry nuclei were both easily detected (Figures 1E and 1F, Movies 1 and 2) and overlapped with the H2B::mCherry nuclear marker (106 dots from 6 gonads traced over time, e.g. Figure 1F). Next, we treated animals with α-amanitin, a Pol II inhibitor that abolishes transcription (Lindell et al., 1970). All MCP::GFP dots disappeared after α-amanitin treatment, and then reappeared after a wash to remove α-amanitin (Figure S1B). Therefore, the MCP::GFP dots reflect transcription.

To ask if Notch-dependent transcriptional activation occurs in bursts, we traced MCP::GFP dots in 3D for several hours and recorded their signal intensities over time (n = 177 dots in 10 gonads). GFP signal intensities oscillated between well above background (“ON”) and indistinguishable from background (“OFF”) (Movies 1 and 2). Because the MCP::GFP dots did not move dramatically within their nucleus (see Methods), we were able to identify individual loci through consecutive bursts.

Importantly, the dynamic MCP::GFP dots were scored at 5 μm intervals across and beyond the GSC pool within the progenitor zone; these 5 μm intervals were then translated to the more traditional measure of number of germ cell diameters (gcd) from the distal end. The vast majority of MCP::GFP dots were restricted to the GSC pool region (1 to ^~^35 μm, or 1 to 6-8 gcd from the distal end), similar to the restriction of *sygl-1* ATS in fixed samples (Lee et al., 2016). The rare MCP::GFP dots found outside that region were just beyond, and no dots were seen proximal to 40 μm or 9 gcd from the distal end. By contrast, nuclei marked with H2B::mCherry were seen throughout the gonad, as expected given the germline promoter (Figure 1E, Movie 1 and 2). We conclude that Notch-dependent activation occurs in bursts within germ cells known to be regulated by GLP-1/Notch signaling from the niche, and that low level or “noisy” bursting was undetectable beyond those cells.

### A gradient in duration of Notch-dependent transcriptional bursts

We next analyzed key dynamic burst features (Figure 2A) for each Notch-dependent burst and did so as a function of position along the gonadal axis (Figure 2B). Transcriptional bursts are episodic with periods of activity (ON-times) punctuated by periods of inactivity (OFF-times). From the intensity of MCP::GFP nuclear signals recorded over time and their normalization to background (see Methods), we determined the duration of ON-time and OFF-time in addition to mean signal intensity over the transcriptionally active period. Each feature was scored at 5 μm intervals across and beyond the GSC pool. Figures 2C and 2D show representative graphs for recordings at two distinct positions: one is near the niche (Figure 2C), comparable to the position of Box 1 in Figure 1E, and one is more proximal (Figure 2D), comparable to the position of Box 2 in Figure 1E.

**Figure 2.**
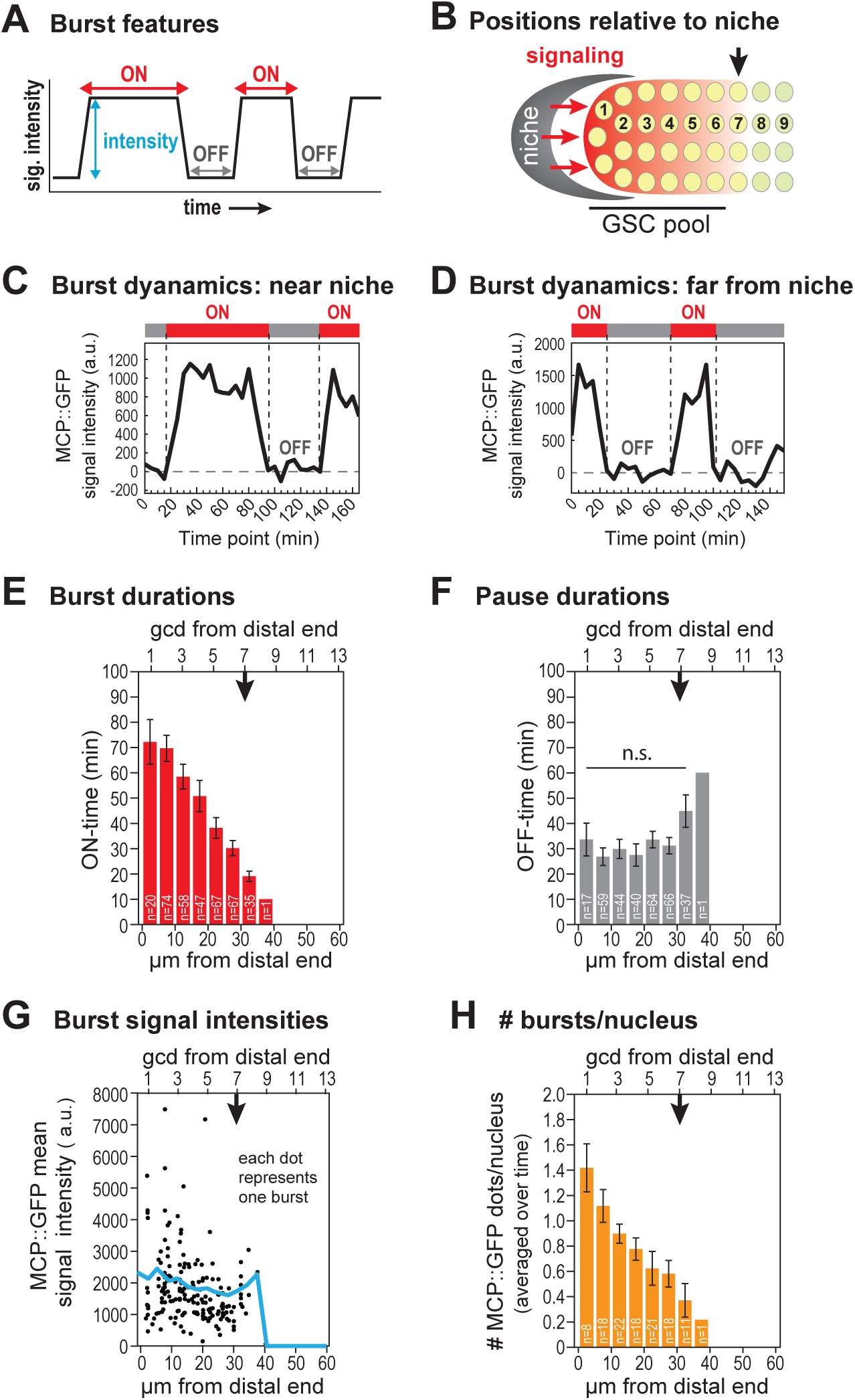
Dynamics of Notch-dependent transcriptional activation. (A) Key features of transcriptional bursts. MCP::GFP signal intensities are recorded over time to determine (1) duration of the burst (ON, red double arrow), (2) duration of the pause between bursts (OFF, grey double arrow) and (3) mean signal intensity during the burst (blue double arrow). (B) Reference for position in the distal gonad. Niche, grey; Notch signaling, red arrows; gradient of Notch transcriptional response, graded red; germ cell nuclei, yellow; digits, number of germ cell diameters (gcd) along the distal-proximal axis from the niche, the convention for germ cell position. The GSC pool extends from gcd 1 to gcd 6-8; the downward black arrow marks the average location where the pool ends (Cinquin et al., 2010; Rosu and Cohen-Fix, 2017). (C,D) MCP::GFP signal intensities (arbitrary unit, a.u.) are traced at 5-minute intervals from a time-lapse movie (Movie 1) and normalized to background (see Methods). Intervals with persisting signals from nascent transcripts (MCP::GFP) are recorded as “ON” (red bar above line plot), and intervals when the signal is essentially at background are recorded as “OFF” (grey bar). (E-H) The downward black arrow marks the average end of the GSC pool, as in 2B. Error bar: standard error of the mean (SEM). (E) Durations of active ON state (ON-times) for all MCP::GFP dots are plotted as a function of position, either in μm (bottom) or number of germ cell diameters [gcd] (top) from the distal end of the gonad. No transcriptional bursts were seen proximal to 40 μm from the distal end within the progenitor zone. n = 460 transcriptional bursts from 177 loci in 10 gonads of living adults. (F) Durations of the inactive OFF state (OFF-times) are plotted as a function of position, as in C. n = 400 rest periods (177 loci from 10 gonads). n.s.: not significant by any pairwise t-test in the bar graph. (G) Signal intensities are averaged over the duration of a transcriptional burst and plotted as a function of position. The blue line marks overall mean signal intensities along the axis, as in C. (H) Number of *sygl-1* active bursts in each nucleus, averaged over time, is plotted as a function of position, as in C.

Our analyses revealed that the lengths of transcriptional burst activity, or ON-times, were sharply graded across the GSC pool -- from ~70 minutes at its distal end to ^~^10 minutes at its proximal end (Figure 2E). By contrast, periods of transcriptional inactivity, or OFF-times, were not graded but instead essentially constant across the pool, with average pauses of roughly half an hour (Figure 2F). The mean intensities of individual bursts were highly variable (Figure 2G, dots), but the averages for all individual bursts at a given position were comparable across the GSC pool (Figure 2G, blue line). However, the number of actively transcribing loci was graded, with the most distal nuclei having the highest average number of MCP::GFP dots (Figure 2H). The overall transcriptional activity per nucleus was therefore also graded when considered at the cellular level (Figure S2A), despite the uniform average burst intensity assessed at the level of individual chromosomal loci (Figure 2G). These burst dynamics are consistent with results from the previously reported smFISH study (Figure S2B). We conclude that the graded transcriptional response to Notch signaling is generated by a gradient in the duration of active transcriptional bursting (ON-time) rather than a gradient in either burst intensity or the duration of transcriptional inactivity (OFF-time). Because ON-times are graded, burst sizes (ON-time x mean signal intensity) are also graded. We conclude that the Notch transcriptional response is “tuned” at the level of burst duration.

### Stochasticity of Notch-dependent burst dynamics

We next investigated the independence or stochasticity of Notch-dependent bursts at individual loci. For this analysis, we first investigated consecutive bursts at the same *sygl-1* locus (Figure 3A). To this end, we not only analyzed data from all consecutive bursts, regardless of position, but also assessed the data as a function of position (Figure 3B, dot colors correspond to position). Essentially no correlation was found for either the summed data (Pearson’s r = 0.1) or position-specific data (r ranged from 0.03 to 0.19, depending on position) (Figure 3B, see legend for r values by position). Moreover, differences in the durations between consecutive bursts were highly variable, ranging over a span of 250 minutes (Figure 3C). We also compared other paired features, such as durations of an active burst and its following inactive pause, durations of a pause and its following burst, and durations of consecutive pauses. These additional pairs also failed to correlate, with r values near zero, regardless of position, and a broad distribution in differences (Figures S3 A-F). Thus, the ON-times of consecutive bursts cannot be predicted even at the same position within the gonad (Figure 3B), despite their gradient (Figure 2E). A similar stochasticity was found when one *sygl-1* locus was compared to a different *sygl-1* locus in the same nucleus (Figure 3D; see Methods). The percentage of time that both loci were in the same state was equivalent to that predicted by chance. We conclude that Notch-dependent ON- and OFF-times are stochastic at any given locus.

**Figure 3.**
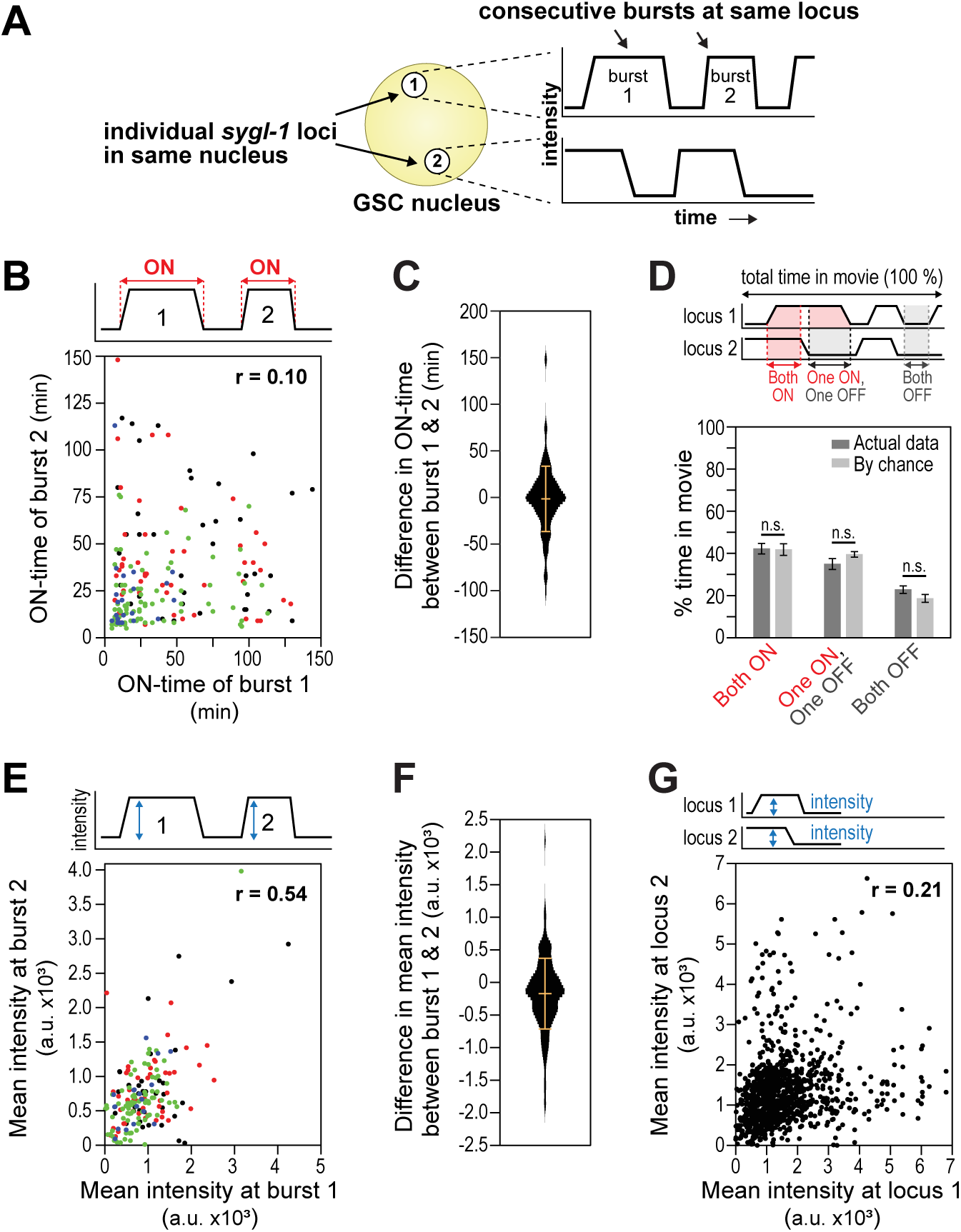
Stochastic Notch-dependent transcriptional dynamics. (A) Experimental design. *Left*, a single germ cell nucleus (yellow) with two *sygl-1* loci (white). *Right*, hypothetical MCP::GFP signal intensities (y-axis) at each locus are plotted over time (x-axis). Comparisons are made in either of two ways – between consecutive bursts at the same locus or between bursts at the two loci. (B) Comparison of burst ON-times for each of two consecutive transcriptional bursts at the same locus. Each dot represents a pair of consecutive bursts (burst 1 & 2). n = 330 pairs. The Pearson’s correlation coefficient for all pairs, regardless of position, shows little or no correlation (r = 0.10); the same is true when analyzed position by position (r-values: 1-10 μm (black dots), −0.13; 10-20 μm (red), −0.16; 20-30 μm (green), 0.19; 30 μm – end (blue), 0.03). (C) Violin plot of differences between ON-times at consecutive burst pairs in B. Bars mark the mean (middle horizontal line, −1.56 minutes) and standard deviation (top and bottom horizontal lines, 35.0 minutes). (D) Comparison of transcriptional states at different *sygl-1* loci in same nucleus over the span of the movie. Both can be active (Both ON), inactive (Both OFF) or distinct (one ON, one OFF). The percentage of time for each situation is plotted. Dark grey bars show data from movies; light grey bars show predictions based on chance (see Methods). Error bar: SEM, n.s.: not significant (*p* > 0.05 by t-test). (E) Comparisons of mean signal intensities for each of two consecutive transcriptional bursts at the same locus. Each dot represents a pair of consecutive bursts (burst 1 & 2). n = 330 pairs. The Pearson’s correlation coefficient for all pairs, regardless of position, shows a modest correlation (r = 0.54); the same is true when analyzed position by position: r-values: 1-10 μm (black dots), 0.54; 10-20 μm (red), 0.46; 20-30 μm (green), 0.63; 30 μm – end (blue), 0.58. (F) Violin plot of differences between mean signal intensities at consecutive burst pairs in E. Bars mark mean (−171.8 a.u.) and standard deviation (541.8 a.u.) as in 3C. (G) Comparisons of mean signal intensities at different loci in the same nucleus over time. n = 1,108 pairs. Pearson’s r = 0.21: little correlation.

Burst intensities, on the other hand, were not fully independent at consecutive bursts from the same locus. Their correlation was modest, either when summed or by position (Figure 3E, r = 0.54 when summed; see legend for r values by position). By contrast, no correlation was seen for random pairings (Figures S4A and S4B) or for pairings between synchronous bursts at different loci within the same nucleus (Figures 3G, S4C, and S4D). In addition, differences between average intensities of consecutive bursts at the same locus varied less than randomly-paired average intensities or those recorded at different loci in the same nucleus (compare Figure 3F to Figure S4B). We conclude that Notch-dependent burst intensities are not fully independent at a single locus, but are independent between loci in the same nucleus.

### Notch NICD modulation of burst dynamics

The graded Notch-dependent transcriptional response reflects a gradient in “signaling strength”, a rough measure of the entire pathway (Lee et al., 2016). To assess the role of the GLP-1/Notch NICD specifically, we employed the temperature-sensitive Notch receptor mutant, *glp-1*(*q224*), which has weaker than normal biological activity at permissive temperature (15°C) and essentially no activity at restrictive temperature (25°) (Austin and Kimble, 1987). This mutant harbors a single amino acid change in its NICD, which weakens stability of the Notch-dependent transcription activation complex (Petcherski and Kimble, 2000). By smFISH, the Notch response was reduced in this mutant at 15°C, including a lower probability of transcription and fewer *sygl-1* nascent transcripts at each ATS (Figure S2B) (Lee et al., 2016). In wild-type animals, by contrast, the probability of transcription and nascent transcript output were both equivalent at 15°, 20° and 25° (Lee et al., 2016).

To understand the role of the NICD in bursting dynamics, we introduced our MS2 system into animals homozygous for *glp-1*(*q224*). The resultant strain was phenotypically indistinguishable from *glp-1*(*q224*) on its own (see Methods). We recorded MCP::GFP signal intensities over time in *glp-1*(*q224*) at 15°C (Movies 3 and 4), and compared their dynamic burst features to wild type (Figure 4). Mutant ON-times decreased roughly three-fold, but, as in the wild type, they were graded across the GSC pool (Figure 4A). Mutant OFF-times, by contrast, increased about 1.5-fold over wild type and were not graded (Figure 4B). As in wild type, the mean intensities for individual bursts were highly variable, but their averages at each position were fairly constant across the GSC pool and about 2-fold lower than wild type (Figure 4C, solid vs dotted line). These decreases in both burst duration and intensity thus lead to a dramatic decrease in burst size (duration x intensity). The number of transcriptionally active loci per nucleus decreased 3-4-fold at each position and were again graded (Figure 4D). These live imaging data are consistent with findings with smFISH (Figure S2B) (Lee et al., 2016). In sum, an NICD that is compromised for assembly into the Notch-dependent transcription activation complex causes decreases in both burst duration and intensity, but as in the wild type, burst duration is graded while intensity is not graded.

**Figure 4.**
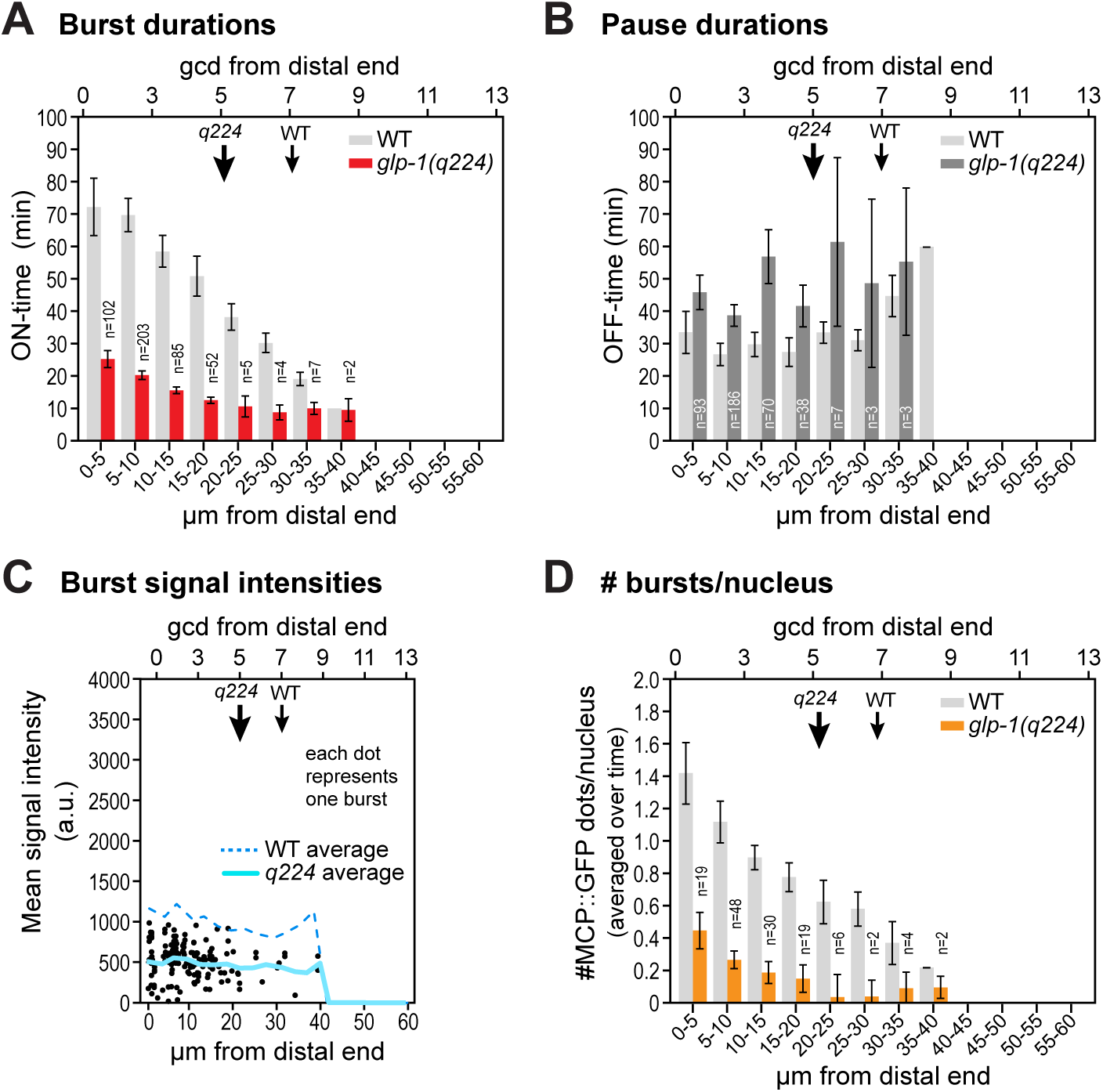
Effect of Notch receptor strength on burst dynamics. (A-D) Comparisons of transcriptional burst features driven by the wild-type (WT) and the weak mutant GLP-1/Notch receptor, *glp-1*(*q224*), all plotted as a function of position. Note that GSC pool size is smaller in the mutant than wild type (downward arrows mark GSC pool end, as in 2B). Error bars: SEM. (A) Burst durations (ON-times) in *glp-1*(*q224*) mutant (red) and WT (light grey). ON-times are dramatically shorter in the mutant, but still graded across the GSC pool. No bursts were seen proximal to 40 μm from the distal end; error bars are omitted with fewer than three bursts. (B) Pause durations (OFF-times) in mutant (dark grey) and WT (light grey). OFF-times are 1.5-fold longer in the mutant than WT, but they remain comparable across the GSC pool (though more variable than WT). (C) MCP::GFP signal intensities as in Figure 2G. All dots reflect data from *glp-1*(*q224*) movies; each dot shows the signal intensity averaged over one burst. Lines mark the average of all these individual mean signal intensities as a function of position: solid light blue line, *glp-1*(*q224*); dashed blue line, wild type. (D) Number of transcriptional bursts per nucleus averaged over time and plotted as a function of position, as in Figure 2H. The mutant receptor lowers that number.

## DISCUSSION

Using the MS2 system and live imaging of intact nematodes, we have visualized Notch-regulated transcriptional bursting over time during normal development. This feat took advantage of a particularly tractable case of Notch signaling that maintains germline stem cells in the nematode *C. elegans*. Our study stands out among other studies of regulated transcriptional bursting by its analysis of regulation in a native metazoan context, and its focus on effects of a canonical signaling pathway.

### Notch-dependent transcriptional activation is “bursty” in its native context

A major conclusion from this work is that Notch-dependent transcription is episodic or “bursty” in intact animals as niche signaling maintains stem cells. Although this was expected, both from the universality of the phenomenon (Chubb et al., 2006; Golding et al., 2005; Raj et al., 2006) and from the probabilistic nature of Notch-dependent transcription seen with smFISH (Lee et al., 2016), other possibilities were feasible. For example, Notch signaling might have driven transcription from an inherently noisy and bursty state to a continuous mode. No studies prior to this work had looked at the dynamics of Notch-dependent transcription, or for that matter any other canonical signaling pathway in its native context. Our results provide compelling evidence that Notch-dependent transcriptional activation is indeed bursty. By extension, we suggest that transcription activated by other canonical signaling pathways will also be bursty *in vivo*.

A growing literature on the regulation of transcriptional bursting in cultured cells has focused on the idea that transcriptional regulators increase burst frequency of otherwise noisy, sporadic transcription (Corrigan and Chubb, 2014; Kafri et al., 2016; Larson et al., 2013; Molina et al., 2013; Senecal et al., 2014). Our analyses of Notch-regulated transcriptional bursting in its native context differ from this consensus in a striking way: *sygl-1* bursting was undetectable outside the region where Notch maintains a stem cell state and prevents differentiation (Austin and Kimble, 1987; Cinquin et al., 2010; Kershner et al., 2014). Why do we not see a low level of “noisy” bursting outside the pool? One part of the explanation is likely detection, because the highly sensitive smFISH did in fact detect exceedingly rare *sygl-1* transcription outside the GSC pool (1 ATS on average per ~130 nuclei in the interval of 50-60 μm from the distal end [11-12 gcd] compared to 96 ATS on average per ~130 nuclei in the interval of 0-10 μm from the distal end [1-2 gcd]) (Lee et al., 2016). But in addition, we suggest that “noisy” transcriptional bursting is likely silenced in its natural *in vivo* setting by other factors or chromatin regulators that drive it to an undetectable level at developmentally key loci. Identification of such regulators is a crucial line of future research.

### Notch signaling strength modulates burst duration

A second major conclusion from this work is that *in vivo* Notch signaling modulates burst duration (ON-time) to determine the probability of its transcriptional activation. smFISH had revealed a spatial gradient of Notch-dependent transcriptional probability (Lee et al., 2016). Here we find that burst ON-times are also steeply graded, much like the transcriptional probability; by contrast, OFF-times and mean burst intensities are not graded but essentially constant across the GSC pool. Therefore, Notch modulation of burst duration appears to be the key determinant of transcriptional probability. This gradient reflects a gradient in Notch signaling strength, but how that strength modulates burst duration remains an open question. The answer is unlikely to depend on stability of the Notch-dependent ternary complex, because that reduces both burst intensity and duration, as seen for the mutant NICD in *glp-1*(*q224*). We suggest therefore that some other aspect of Notch signaling is likely graded, such as ligand concentration. Regardless, this system can now be used to investigate how burst duration is regulated in an *in vivo* setting. Questions include whether burst duration is determined by the turnover dynamics of chromatin modifications or how long the promoter is sustained in a phase-separated state.

The Notch tuning of burst duration differs markedly from what has been found for other metazoan transcriptional regulators. For example, steroid-mediated gene activation increased burst frequency by shortening burst pauses (Fritzsch et al., 2018; Larson et al., 2013), and Wnt signaling increased burst frequency by modulating both ON- and OFF-times (Kafri et al., 2016). Indeed, modulation of burst frequency via regulation of OFF-times has been suggested as a universal phenomenon (Nicolas et al., 2017). We consider two possible explanations for our distinct results. First, the Notch effect on burst duration may be a special feature of Notch regulation. Indeed, a strikingly similar effect of Notch signaling on burst duration was discovered independently in *Drosophila* embryos (Falo-Sanjuan et al., accompanying mss.), suggesting that it is a conserved phenomenon. Second, the Notch effect on burst duration may represent regulation in a native context, which was used for both our study and that in *Drosophila* (Falo-Sanjuan et al., accompanying mss.), but is rare among other studies. To distinguish between these possibilities, the effects of other canonical signaling pathways must be analyzed, and if possible analyzed in their native context.

### Effect of Notch NICD strength and stability of the Notch transcriptional complex

Most analyses in this work were done with the wild-type GLP-1/Notch receptor in its native context (with exception of the added MS2 system, which had no detectable phenotypic effect). But we also examined a mutant GLP-1/Notch receptor, again in its native context. This mutant harbored an amino acid substitution in the NICD, which led to weaker than normal assembly into the Notch-dependent transcription factor complex (Petcherski and Kimble, 2000). The weaker NICD reduced burst ON-time and mean burst intensity, both by 2-3 fold, and increased OFF-time by ~1.5 fold. Thus, stability of the transcription factor complex dramatically reduces burst size, which is a function of both burst duration and burst intensity.

The difference between the broad effect of this weak NICD on multiple burst features and the more specific effect of a graded signaling on burst duration is striking. One might have thought, *a priori*, that the readout would have been more similar. Yet the differences are intriguing and lead to many questions. Do individual components of the Notch pathway affects burst dynamics in distinct ways? Will all components of the transcription factor complex behave like the NICD or will they have individual roles? Will distinct Notch ligands direct specific burst behaviors, as suggested recently, albeit with a vastly different assay and kinetics (Nandagopal et al., 2018)? Do chromatin modifiers affect the NICD readout differently than the wild-type gradient readout? Do intrinsically disordered domains in pivotal components of the transcription factor complex affect one readout over the other? Now that Notch-dependent transcriptional bursting can be assayed in a natural setting, these questions can be addressed rigorously with potential impact for identifying how to target components for manipulation in humans.

### Stochasticity of Notch-dependent ON-times and OFF-times

The stochasticity of Notch-dependent transcription was first discovered with smFISH (Lee et al., 2016), but the current live imaging analysis clarified its stochasticity in terms of key burst features. Thus, the ON- and OFF-times are not correlated with each other, either for consecutive bursts at the same locus or for bursts at distinct loci within the same nucleus (r ≤ 0.10 for the various pairings). The one exception is mean burst intensity, which shows a modest correlation (r = 0.54) for consecutive bursts at the same locus. Because no correlation was seen for bursts at distinct loci in the same nucleus, either by smFISH or live imaging, we suggest that individual promoters may adopt a “configuration” that is sustained, at least in part, for consecutive bursts. That configuration might involve, for example, promoter-specific chromatin modifications (e.g. Lenhard et al., 2012) or promoter-specific phase-separation (e.g. Hnisz et al., 2017). Yet the more striking result is the lack of correlation between most burst features.

## Supporting information

## ACKNOWLEDGEMENTS

We thank Tina Lynch and Sarah Crittenden for critical comments on the manuscript. We also thank Anne Helsley-Marchbanks for help preparing the manuscript and Laura Vanderploeg for help with the figures. The codon-optimized superfolder GFP was a gift from Andy Golden and Harold Smith. This work was supported in part by the American Heart Association (18POST34030263) to CL. JK is an Investigator of the Howard Hughes Medical Institute.

## AUTHOR CONTRIBUTIONS

CL and JK designed experiments and oversaw the project. CL and HS adapted the MS2 system for this project. CL adapted methods for live imaging to this project, generated movies and wrote software to analyze images. CL and JK wrote the paper with input from HS.

## Methods

### KEY RESOURCE TABLE

| Reagent or Resource | Source | Identifier |
| --- | --- | --- |
| **Experimental models: Organisms/Strains** |  |  |
| Wild-type *C. elegans* | *C. elegans* Genetics Center (CGC) | N2 Bristol |
| *ttTi5605 II; unc-119(ed3) III* | Frøkjær-Jensen et al. (2008) | EG6699 |
| *unc-119(ed3) III; oxTi365 V* | Frøkjær-Jensen et al. (2014) | EG8082 |
| *glp-1(q224ts) III* | Kodoyianni et al. (1992) | JK4605 |
| *sygl-1(q828) I; qSi49[Psygl-1::3xFLAG::sygl-1::sygl-1 3'end] II* | Shin et al. (2017) | JK5499 |
| *qSi369 [Psygl-1::24xMS2 loops::3xFLAG::sygl-1::sygl-1 3'end] II; unc-119(ed3) III; qSi370[Pmex-5::MS2 coat protein::linker::GFP::tbb-2 3'end::gpd-2 intergenic sequence::H2B::mCherry::unc-54 3'end] V* | This work | JK5896 |
| *qSi369[Psygl-1::24xMS2 loops::3xFLAG::sygl-1::sygl-1 3'end] II; glp-1(q224) III/hT2[qIs48](I,III); qSi370[Pmex-5::MS2 coat protein::linker::GFP::tbb-2 3'end::gpd-2 intergenic sequence::H2B::mCherry::unc-54 3'end] V* | This work | JK5943 |
| *sygl-1(q828) I; qSi369[Psygl-1:: 24xMS2 loops::3xFLAG::sygl-1::sygl-1 3'end] II; qSi370[Pmex-5::MS2 coat protein::linker::GFP::tbb-2 3'end::gpd-2 intergenic sequence::H2B::mCherry::unc-54 3'end] V* | This work | JK5932 |
| **Antibodies** |  |  |
| Mouse α-FLAG | Sigma | F1804 |
| Donkey α-Mouse IgG (H+L), Alexa Fluor 647 | Invitrogen | A31571 |
| **Plasmids** |  |  |
| *Psygl-1::24xMS2 loops::3xFLAG::sygl-1::sygl-1 3'end*, cloned into pCFJ151 | This work | pJK2014 |
| *Pmex-5::MS2 coat protein::linker::GFP::tbb-2 3'end::gpd-2 intergenic sequence::H2B::mCherry::unc-54 3'end*, cloned into pCFJ151 | This work | pJK2020 |
| **Software and algorithms** |  |  |
| MATLAB | Mathworks | R2018b |
| FIJI | ImageJ | 1.52h |
| LAS image acquisition software | Leica Biosystems | 3.3.1 or X |
| Micromanager | www.micro-manager.org | 1.4 |
| Custom MALTALAB code | This work<br>github.com/chlasic/MS2_analysis | N/A |
| Image J Plugin, StackReg | imagej.net/StackReg | N/A |
| **Reagents for live imaging** |  |  |
| CherryTemp | Cherry Biotech, Rennes, France | N/A |
| Polystyrene microbeads, 0.1 μm diameter | Polysciences | 00876 |
| α-amanitin | Sigma | A2263-1MG |
| Serotonin | Sigma | H7752-5G |

### CONTACT FOR REAGENT AND RESOURCE SHARING

Further information and requests for resources and reagents should be directed to and will be fulfilled by the corresponding author, Judith Kimble.

### EXPERIMENTAL MODEL AND SUBJECT DETAILS

#### Nematode strains and culture

Most strains were maintained at 20°C as described in (Brenner, 1974); those carrying *glp-1*(*q224*) (Austin and Kimble, 1987) were maintained at 15°C. The wild type was N2 Bristol. Alleles, transgenes, and balancers are as follows: LG I: *sygl-1*(*q828*) (Shin et al., 2017), *hT2[qIs48]* (Siegfried and Kimble, 2002). LG II: *ttTi5605* (Frøkjær-Jensen et al., 2008), *qSi49[Psygl-1::3xFLAG::sygl-1::sygl-1 3’end]* (Shin et al., 2017), *qSi369[Psygl-1::24xMS2 loops::3xFLAG::sygl-1::sygl-1 3’end] II* (this work). LG III: *glp-1*(*q224*) (Kodoyianni et al., 1992). LG *V: oxTi365* (Frøkjær-Jensen et al., 2014), *qsi370[Pmex-5::MS2 coat protein::linker::GFP::tbb-2 3’end::gpd-2 intergenic sequence::H2B::mCherry::unc-54 3’end]* (this work).

#### Generation of strains carrying an MS2 system in the *C. elegans* germline

Constructs: (1) pJK2014 *[Psygl-1:: 24xMS2 loops::3xFLAG::sygl-1::sygl-1 3’end]*. The construct containing *sygl-1* tagged with 24xMS2 loops was cloned in two steps. First, the *sygl-1* gene, including its open reading frame and previously described 5’ upstream and 3’ downstream sequences (Shin et al., 2017), was cloned into the *Spe* I site of pCFJ151 UniMos vector (Frøkjær-Jensen et al., 2008) using the Gibson assembly method (Gibson et al., 2009). At this step, *Not* I and *Pme* I restriction sites were inserted just in front of the start codon of *sygl-1*. Second, to insert 24xMS2 loops in front of the start codon, the aforementioned plasmid and pCR4-MS2 (Addgene #31865) (Bertrand et al., 1998) were digested with *Not* I and *Pme* I restriction enzymes and ligated together using T4 DNA ligase (Roche #10481220001).

The final product is pJK2014. (2) pJK2020 *[Pmex-5::MS2 coat protein::linker::GFP::tbb-2 3’end::gpd-2 intergenic sequence::H2B::mCherry::unc-54 3’end]*. This construct contains MS2 coat protein tagged with a codon optimized superfolder GFP (MCP::GFP) and mCherry-tagged histone 2B (H2B::mCherry), driven by the *mex-5* promoter and *tbb-2* (for MCP::GFP) and *unc-54* (for H2B::mCherry) 3’end regulatory sequences (Merritt et al., 2008); MS2 coat protein (Addgene #27121) was fused with superfolder GFP (Kersey et al., 2016); Histone 2B was fused with mCherry (Merritt et al., 2008); *gpd-2* intergenic sequence was used to generate a trans-spliced operon (Huang et al., 2001). Of note, the nuclear localization sequence (NLS) in the original construct (Addgene #27121) (Fusco et al., 2003) was removed to minimize the MCP::GFP background in the nucleus. All amplified fragments were targeted to the *Spe* I site of the pCFJ151 UniMos vector (Frøkjær-Jensen et al., 2008) using the Gibson assembly method (Gibson et al., 2009). The final product is pJK2020.

Transgenes: Single-copy transgenes were generated using the Mos1-mediated single-copy insertion method (MosSCI) (Frøkjær-Jensen et al., 2008). Briefly, 50 ng/μl plasmid (pJK2014 or pJK2020, see Constructs) was microinjected into EG6699 or EG8082 along with transposase and co-injection markers. Integration was screened by Unc movement rescue, and further confirmed by PCR amplification of Mos insertion sites. At least two lines were generated for each construct and representative lines were selected for further characterization: *qSi369* for *24xMS2 loops::sygl-1* and *qSi370* for *MCP::GFP; H2B::mCherry*.

Final strains: To generate JK5896, a strain with the complete MS2 system in an otherwise wild-type background, animals carrying *qSi369* or *qSi370* were crossed with each other, and their progeny were validated using PCR and microscopy (green dots for *qSi369* and red germ cell nuclei for *qSi370*). JK5896 had a normal growth rate (^~^4 days to become adult, n=75), brood size (^~^270 progeny after becoming gravid, n=5) and progenitor zone size (~20 gcd, n=21) at 20°C, similar to those previously reported for wild type (Crittenden et al., 2006; Muschiol et al., 2009; Nehammer et al., 2015). To generate JK5943, a strain with the complete MS2 system in a *glp-1*(*q224*) background, JK5896 was crossed to *glp-1*(*q224*) (JK4605). A progeny of this cross was mated into the genetic balancer *hT2[qIs48]* for strain propagation purposes, and each locus was further homozygosed. The final progeny was validated by PCR, microscopy, and sterility at 25°C. Only JK5943 heterozygous for *glp-1*(*q224*), which was balanced with *hT2[qIs48]*, was fertile at 20°C. For experiments, non-balancer carrying progeny of JK5943 homozygous for the *glp-1*(*q224*) allele were used. These animals raised at 15°C behaved comparably to JK4605 with respect to growth rate (~5 days to become adult, n=75) and progenitor zone size (~10 gcd, n=21), as previously reported for JK4605 (Austin and Kimble, 1987; Fox and Schedl, 2015; Lee et al., 2016).

#### Worm immobilization for long-term live imaging

Microscope slides for long-term live imaging were prepared as previously described (Kim et al., 2013), with a few modifications. First, 5-7.5% (w/v) agarose gel-pad slides (dissolved in M9) were freshly prepared. Then, a 0.5-2 μL suspension of polystyrene microbeads (Polysciences, 2.5% by volume, 0.1 μm diameter) was added to the middle of the gel-pad, immediately followed by serotonin (final concentration 20-25 mM, diluted in M9) to create a mixture of microbeads and serotonin. This mixture efficiently immobilizes worms on the microscope slide without affecting or halting their pharyngeal movement (3-5/sec), egg laying (2-5/hrs) or germ cell division (6.25 M-phases seen on average within any one hour in the region of 1 to 7-8 gcd), consistent with previous reports (Rosu and Cohen-Fix, 2017; Song and Avery, 2012; Teshiba et al., 2016) (also see Results). 10-15 staged young adult worms (24h past mid-fourth larval [L4] stage) were transferred to the gel-pad before the serotonin and microbeads solution dried (within a minute). A cover slip was placed gently on the gel-pad. VALAP (Vaseline, lanolin, and paraffin) was applied around the edges of the coverslip to prevent the microscope slide from drying over time. The slide was attached to a CherryTemp chip (Cherry Biotech, Rennes, France) and mounted on the CherryTemp chip holder located on the confocal microscope stage to keep a consistent temperature throughout the live imaging process. Live imaging was performed immediately afterwards.

#### Confocal microscopy setup for MS2 live imaging

All imaging was done using a Leica TCS SP8 (confocal laser scanner) equipped with a Leica HC PL APO CS2 63x/1.40 NA oil immersion objective, two sensitive hybrid detectors (HyDs) and standard and LAS image acquisition software (version 3.3.1 or X, Leica Microsystems Inc., Buffalo Grove, IL). Two channels were imaged simultaneously to capture MCP::GFP and H2B::mCherry at the same time, with bidirectional scanning at 900 Hz and 250% zoom factor with 512X512 or 1024X512 resolution. The green channel was imaged with the excitation laser at 488 nm (0.3-0.4% laser power, Argon, 40% gain) and the red channel with the excitation laser at 594 nm (0.6-1% laser power, HeNe, 40% gain), with a pinhole size at 105.1 μm. A line average of two snapshots was used for all channels. All imaging was done with HyDs, including DIC. Signal acquisition windows (a range of wavelength in which signals are collected by HyDs) were carefully selected to minimize bleed-through (each window started 10-nm longer than the excitation laser and spanned 50 nm). All gonads were imaged with a total z-depth of >15 μm and a z-step size of 0.4 μm. Worms that were not completely immobilized on the slide, or whose pharynges were not actively pumping, were excluded from live imaging and analyses. Multi-point imaging and autofocusing functions embedded in the LAS image acquisition software (3.3.1 or X) were used during all image acquisitions. Up to six gonads were imaged in each set of time-lapse recordings. Images were initially taken every 1, 2, 5, or 10 minutes, and a 5-minute interval was chosen as an optimal setting to capture the dynamics of the MCP::GFP signals with minimal light exposure. Autofocusing was conducted at every other time point in the DIC channel using the 594-nm laser. Occasionally, gonadal image drifting caused by slight movements of the animal or its gonad was corrected manually. For all imaging, the temperature controller, CherryTemp (Cherry Biotech, Rennes, France) with its accompanying software (Cherry Biotech TC) was used (wild type at 20°C, *glp-1*(*q224*) at 15°C).

#### Image processing and analysis

Images of *C. elegans* gonads in individual MS2 movies were aligned in two steps using customized, automated ImageJ (version 1.52h) macros (see Data Availability). These macros use the ImageJ plug-in “StackReg” (Thevenaz et al., 1998) with modifications (e.g. choice of the reference image, direction for the alignment). First, gonads were aligned along the z-axis at each time point using the middle plane of the z-stack (the thickest region in the gonad) as a reference image. Then, these aligned z-stacks (at each time point) were aligned again through all time points to keep the gonad in the same position throughout the MS2 movies. These processes correct for subtle movements of the worm or natural gonadal displacements due to intestinal movement during feeding. A few images that were not properly corrected by the automated ImageJ macros were manually aligned, using customized ImageJ codes (see Data Availability). All gonadal images were split in halves on the z-axis for further analysis to minimize overlap of germ cell nuclei in the z-projected images. A circular region of interest (ROI, 1 μm diameter) was drawn on each MCP::GFP dot in the z-projected (sum slices) gonadal images to measure signal intensity. The same ROI was used at all time points to record intensities over time. To measure the background signal intensity, at least three of the ROIs (1 μm diameter) were randomly drawn in nuclei for each image and each time point, and their intensities were recorded separately. For normalization, the mean background intensity was subtracted from raw MCP::GFP intensities. Further analyses used the normalized signal intensities.

Most MCP::GFP dots did not move dramatically inside the nucleus (95% (57/60) move <1 μm). All MCP::GFP dots that moved less than 1.5 μm (150% of ROI) between two consecutive time points (5 minutes apart) were considered as an MCP::GFP signal from the same locus. Each ROI position was kept in the same coordinates within the nucleus when the tracked MCP::GFP dot disappeared during the pause between active transcriptional bursts until it reappeared. The duration of transcriptional bursting (ON-time, its intensity, OFF-time) and all other measurements (e.g. plots in Figure 3) were calculated and generated using customized MATLAB codes (see Data Availability). The beginning and the ending of each transcriptional event (e.g. bursting) were defined similarly to other previously reported methods (Corrigan et al., 2016; Larson et al., 2013). Specifically, ON-time was measured only when raw MCP::GFP dot intensity was sustained at least 50% higher than the raw background. OFF-time was scored for time duration between two bursts. To quantitate the mean of burst signal intensities, the normalized MCP::GFP signal intensities (after background subtraction) during each transcriptional burst were summed over all time points during the burst, then each was divided by the length of ON-time of the corresponding burst. To quantitate the number (Figure 2H) or summed intensity (Figure S2A) of MCP::GFP dots per nucleus, each nucleus was scored for the number of MCP::GFP dots inside the nucleus as well as their summed signal intensity for each time point throughout the movie. Then the measured data (# dots or intensities) were averaged over time for each nucleus.

To compare transcriptional states at two different loci in the same nucleus over the span of the movies (Figure 3D), the probability of a transcriptional state of two paired chromosomal loci (Both ON; One ON, one OFF; or Both OFF) was estimated by the product of individual probabilities of transcriptional state at each locus (e.g. “probability of ON state at locus 1” X “probability of ON state at locus 2” to estimate the probability of ‘Both ON’). The probability of ON state at one locus was estimated from the percentage of time when that locus was transcriptionally active (MCP::GFP signal is on) during the whole movie (time period of ‘ON state’ divided by the total length of the whole movie). In contrast, the probability of one locus being inactive (OFF state) is the ratio of the sum of all OFF-times divided by the length of the whole movie. To estimate the probability of ‘Both ON’ at paired two loci in the nucleus, two probabilities of the ON state calculated for each locus were multiplied. The probabilities of transcriptional state at two loci (Both ON; One ON, one OFF, or Both OFF) were then converted to percentages by multiplying by 100 (Figure 3D, ‘by chance’), and compared with the percentage of time measured from the MS2 movies (Figure 3D, ‘actual data’). The comparison between the probability (by chance) and measurement (actual data) allows us to test whether the transcriptional states of two loci in the same nucleus are dependent or not. In other words, this statistical test distinguishes whether the overlap of bursts observed in the actual data (both ON) is merely by chance. If the actual data for ‘both ON’ are higher than its probability ‘by chance’, transcriptional burst at one locus may promote the burst at the other locus. The opposite case suggests a mutually exclusive transcriptional activation at different loci in the same nucleus.

For figure preparation, all images were processed with linear contrast enhancement in ImageJ (version 1.52h) using a minimum contrast of 1.10X mean background signal intensity and a maximum contrast of 1.25X maximum signal intensity as used in a previous study (Lee et al., 2017; Lee et al., 2016). Customized MATLAB codes (see Data Availability) were used for generating the plots used in the figures.

#### Immunostaining and DAPI staining

Immunostaining followed established protocols (Crittenden et al., 2017). Briefly, dissected gonads were fixed with 3% paraformaldehyde for 30 minutes and permeabilized with ice-cold methanol for 10 minutes. Next, samples were incubated with anti-FLAG primary antibody overnight [1:1000, Sigma #F1804)], followed by a 1-hour incubation with secondary antibody [donkey Alexa 647 anti-mouse (1:500, Invitrogen #A31571)] at room temperature. DAPI was added at a final concentration of 0.5–1 ng/μl during the last 10 minutes of the secondary antibody incubation. Vectashield (Vector Laboratories #H-1000) was the mounting medium.

#### Compound Microscopy

A Zeiss Axioskop equipped with a 63×1.4NA Plan Apochromat oil immersion objective and an ORCA cMOS camera was used. Carl Zeiss filter sets 49, and 43HE were used for the visualization of DAPI and Alexa 647 respectively. An X-Cite 120Q lamp (Lumen Dynamics) was the fluorescence light source.

#### α-amanitin treatment

α-amanitin treatment was performed as previously described (Lee et al., 2017; Lee et al., 2016). Briefly, staged young adults (24h post mid-L4 stage) were treated with α-amanitin dissolved in M9, at 100 μg/mL for 1.5 hrs at 20°C on a Shaker rotisserie (Thermo Scientific, ~0.85 rad/s). Following treatment, worms were washed three times with M9. Images were taken before treatment, after 1.5-hr treatment and 1 hr after the washes.

### QUANTIFICATION AND STATISTICAL ANALYSIS DATA

Sample sizes are indicated in the figure legends. Error bars indicate the standard error of the mean (SEM) unless noted otherwise. To calculate the statistical significance between two groups, Student’s (for groups with equal variance) or Welch’s (for groups with unequal variance) two sample t-test was used after the normality test using quantile plots, unless noted otherwise. For comparing multiple groups, an ANOVA was used first and then the pairwise comparisons followed (only *p*-values were reported). *p*-values smaller than 0.01 were considered to be a significant difference. To test correlation between two variables, Pearson’s correlation coefficient (Pearson’s r) was calculated. 0 ≤ |r| < 0.20 was considered as no or little correlation; 0.20 ≤ |r| < 0.40: weak (positive or negative) correlation; 0.40 ≤ |r| < 0.60: modest correlation; 0.60 ≤ |r| < 0.80: strong correlation; and 0.80 ≤ |r | ≤ 1: very strong correlation. See ‘Image processing and analysis’ for quantification of transcriptional dynamics in MS2 movies.

### DATA AVAILABILITY

All customized MATLAB codes and ImageJ macros used in this study are deposited and are available at: https://github.com/chlasic/MS2_analysis.

Several ImageJ macros use the plug-in ‘StackReg’ (https://imagej.net/StackReg), which must be installed in the ImageJ ‘plugins’ folder prior to using the customized macros. For automated MS2 movie alignments, two ImageJ macros were used sequentially: ‘MultiStackReg_BatchProcess1of2.ijm’ and ‘MultiStackReg_BatchProcess2of2.ijm’. The former code aligns gonadal images through the z-axis and the latter does so through the time points. In cases where no DIC channels were included in the z-stacks, ‘MultiStackReg_BatchProcess1of2_2Ch.ijm’ and ‘MultiStackReg_BatchProcess2of2_2Ch.ijm’ were used instead. For manual MS2 movie alignments, either one of following ImageJ macros can be used: ‘TranslateStack.ijm’ or ‘TranslateStackAllAfterPos.ijm’. The former code aligns the image only at the current time point, and the latter does for whole time points after the current time point.

To analyze the MCP::GFP tracking data from MS2 movies, a custom MATLAB code ‘MS2track_analysis.m’ was used. This code includes multiple customized functions to analyze all features of transcriptional dynamics quantitated in this study (e.g. burst duration, intensity). This code requires other customized MATLAB functions ‘MS2normalize.m’ and ‘MS2sumInt.m’.

### Supplemental figure legends

**Figure S1.**
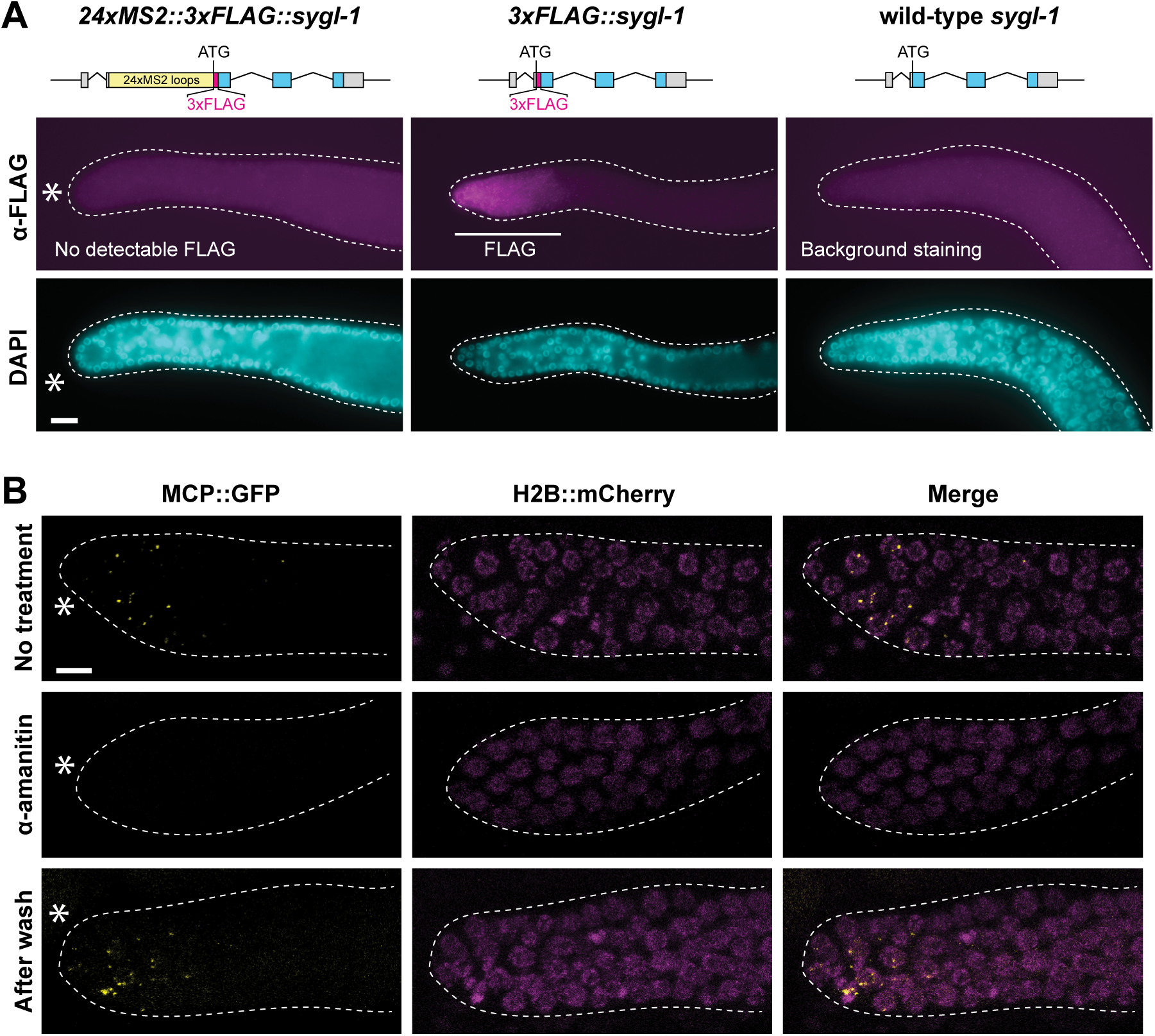
MCP::GFP as a direct reporter of *sygl-1* Notch-dependent transcriptional activation. (A) Immunostaining of dissected adult gonads to see FLAG epitope tag fused to SYGL-1. Schematic of each transgene is shown above its respective confocal images. DAPI visualizes germ cell nuclei. Asterisk, distal end. Scale bar, 10 μm. (B) *Top, sygl-1* nascent transcripts are seen as bright dots in the distal gonad using the MS2 system, as in Figure 1E. *Middle*, no bright dots are seen after 1.5 h treatment with 100 μg/mL α-amanitin, which blocks transcription. *Bottom*, bright dots are restored after α-amanitin is washed out. Dashed line, gonadal outline; asterisk, distal end. Scale bar, 5 μm.

**Figure S2.**
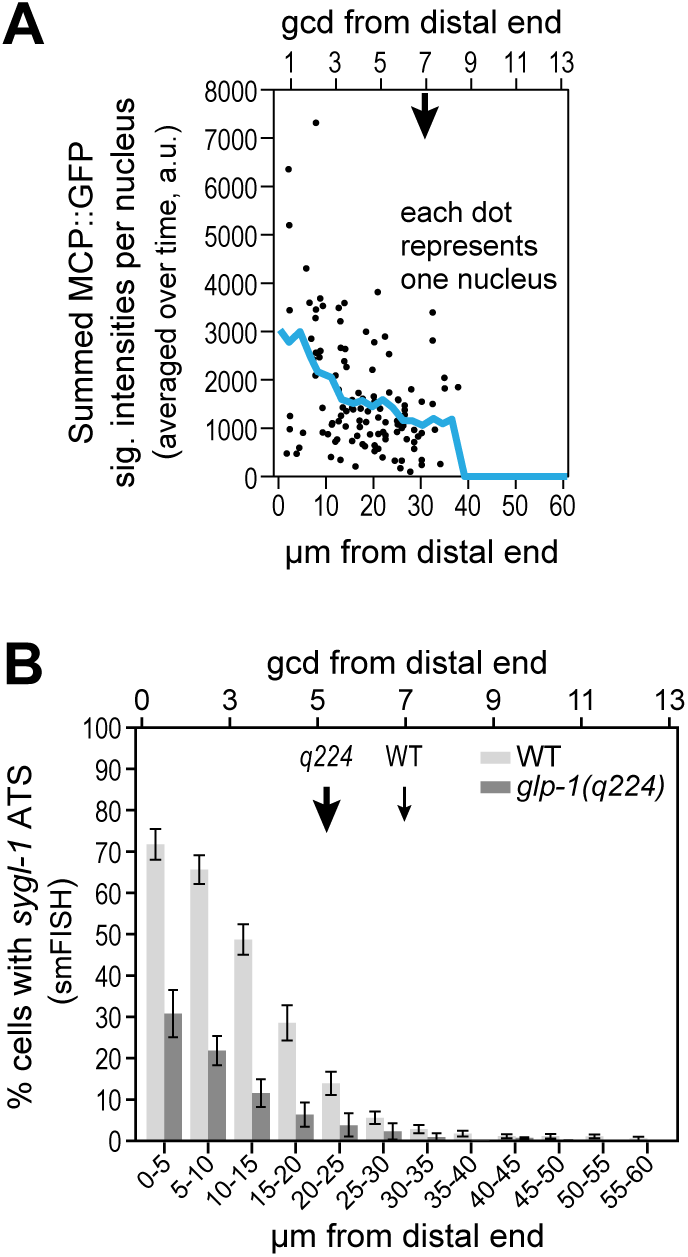
Notch-dependent transcriptional dynamics is graded. (A) Signal intensities from MCP::GFP dots were averaged across each burst and then summed within each nucleus. These summed MCP::GFP signal intensities per nucleus are plotted as a function of position, as in Figure 2E. The blue line marks the average of all intensities from individual nuclei as a function of position. The downward arrow marks the proximal boundary of the GSC pool, as in Figure 2B. (B) smFISH data from Lee et al. (2016) for comparison to live imaging data. This graph shows the percentage of germ cells with at least one *sygl-1* active transcription site (ATS) as a function of position in wild type (light grey) and *glp-1*(*q224*) mutants (dark grey). Downward arrows mark proximal boundary of GSC pool, as in Figure 2B. Error bar: SEM.

**Figure S3.**
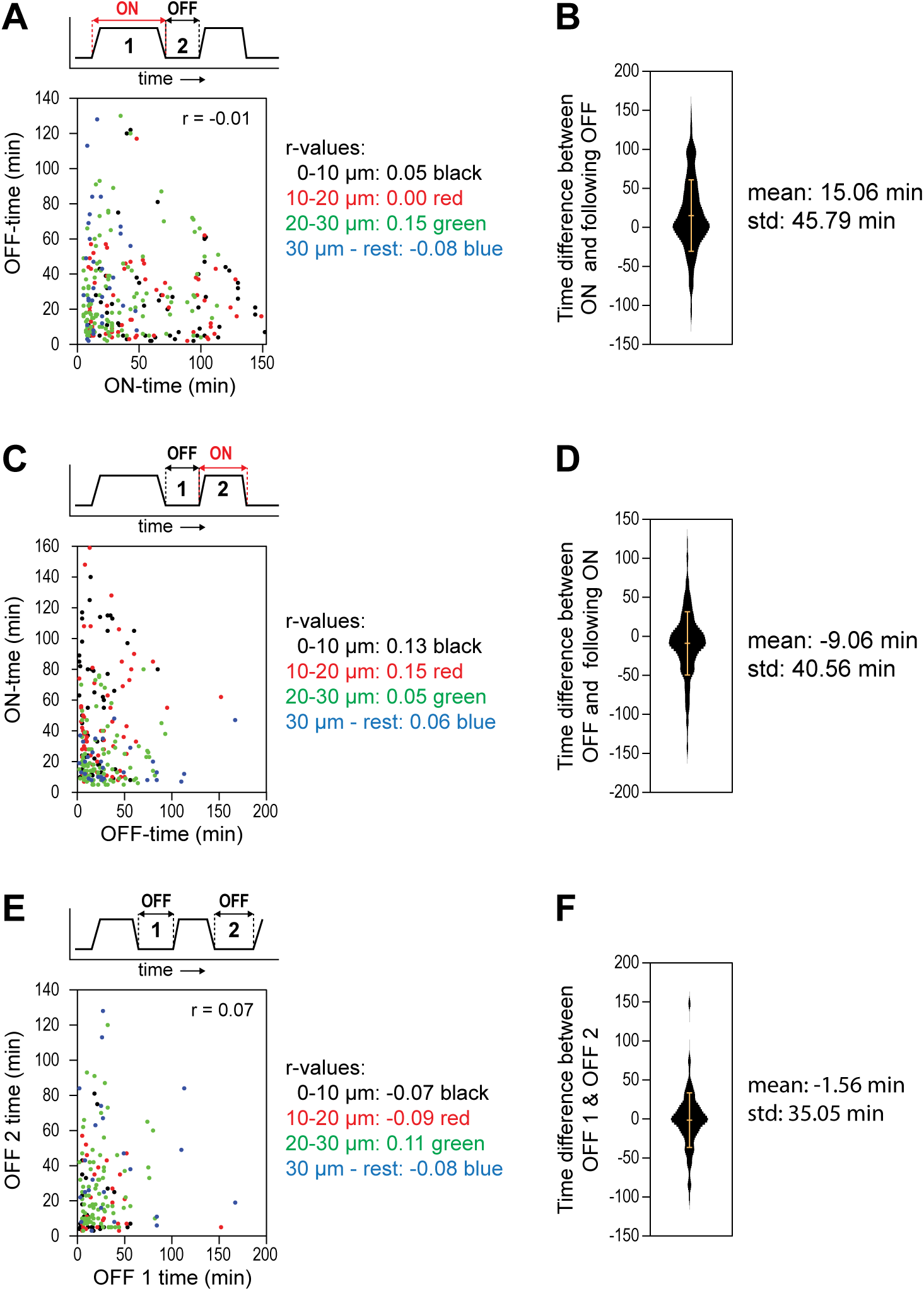
Stochasticity of Notch-dependent transcriptional dynamics. (A, C, E) Each dot represents a pair of consecutive transcriptional states, diagrammed above each plot. n > 428 pairs for all plots. Pearson’s r close to zero: no correlation. (A) Comparison of the duration of an active burst (ON) with the duration of its following inactive pause (OFF) at the same locus. (C) Comparison of the duration of a pause and its following burst at the same locus. (E) Comparison of the durations of consecutive pauses. (B,D,F) Duration differences between pairs of consecutive transcriptional states in A,C, or E, respectively. Bars mark the mean (middle horizontal line) and standard deviation (top and bottom horizontal lines).

**Figure S4.**
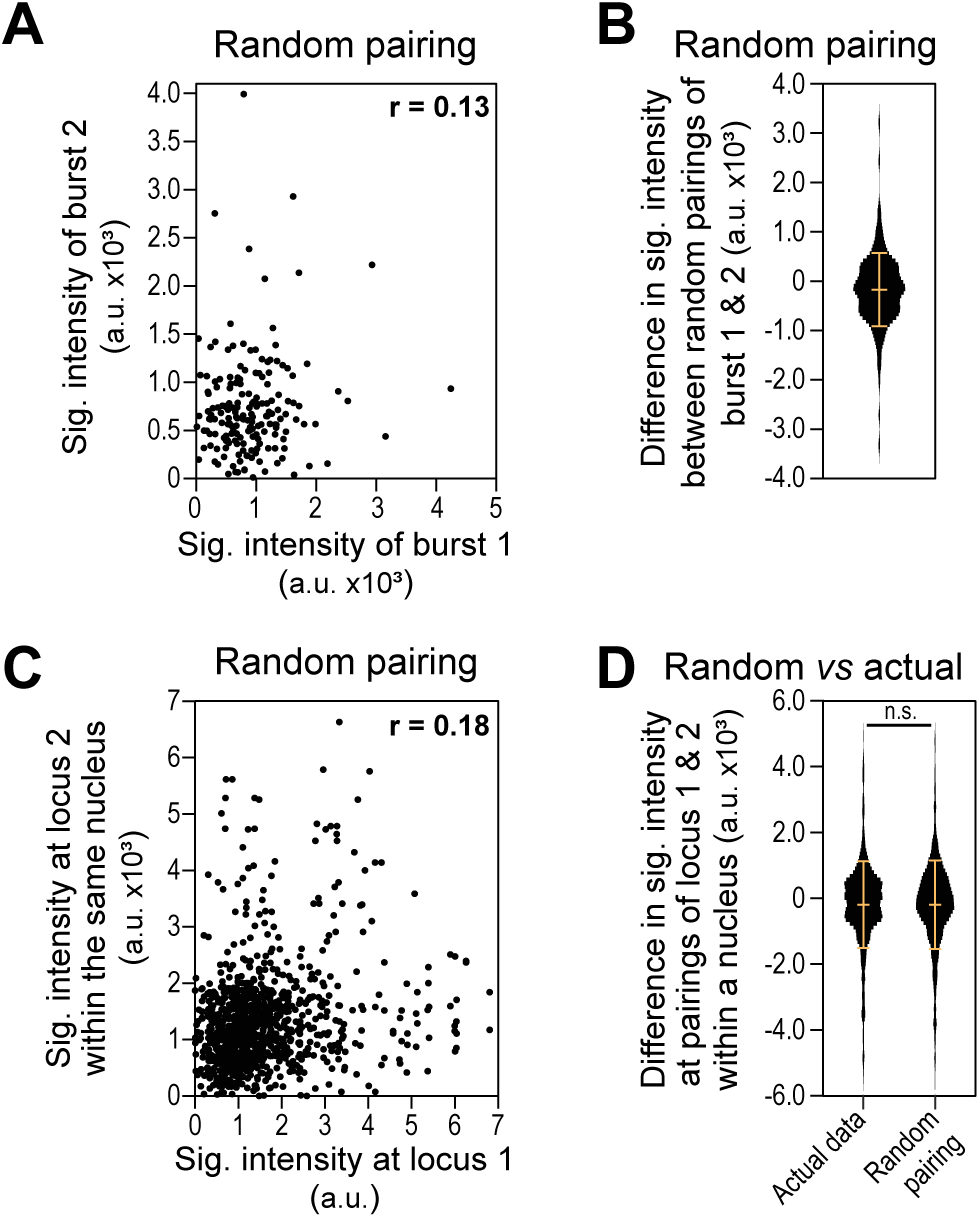
Random pairing simulation of Notch-dependent transcriptional activation. (A) Comparisons of mean signal intensities between two randomly selected transcriptional burst pairs. Data from Figure 3E were used for random pairing. n = 330 pairs. Pearson’s r = 0.13: no correlation. (B) Violin plot of the difference in mean signal intensity between randomly selected burst pairs in A. Bars mark mean (−171.8 a.u.) and standard deviation (741.35 a.u.). (C) Comparison of mean signal intensities at two randomly paired loci. Data from Figure 3G were used for random pairing. n = 1,108 pairs. Pearson’s r = 0.18: no correlation. (D) Violin plot of the difference in mean signal intensity between consecutive burst pairs from Figure 3G (Actual data) or random loci pairs from Figure S4C (Random pairing). Bars mark mean and standard deviation. n.s.: not significant (*p* > 0.05 by t-test).

**Movie 1-4.** Live imaging of Notch-dependent transcriptional activation. MS2 system visualizes *sygl-1* nascent transcripts in germ cell nuclei in the gonad of a live *C. elegans*. The movies focus on the distal region of the gonad (distal end is located at the left in the movie). Bright yellow nuclear dots show *sygl-1* transcripts generated at the ATS using MS2 system, and magenta donut-shaped circles show germ cell nuclei by H2B::mCherry. Cytoplasmic MCP::GFP constitutes background throughout the gonad (yellow haze in cytoplasm) but *sygl-1* nascent transcripts are distinguishable due to relatively low background in the nucleus and bright signal from nuclear MCP::GFP dots. Images were taken every 5 minutes for several hours in wild type (Movies 1 and 2) or *glp-1*(*q224*) mutants (Movies 3 and 4). All movies show z-projection of z-stack images with z-step depth of 0.4 μm.

